# Next Generation AAV-F Capsid gene therapy rescues disease pathology in a model of Pyruvate Dehydrogenase Complex Deficiency

**DOI:** 10.1101/2025.07.30.667478

**Authors:** Anna Keegan, Ozge Cetin, Ellie M Chilcott, Juan Antinao Diaz, Simon Eaton, Simon N Waddington, John R Counsell, Shamima Rahman, Rajvinder Karda

**Affiliations:** EGA-Institute for Women’s Health, University College London, London, UK; Unit of Paediatric Surgery, UCL Institute of Child Health, London, UK; Research Department of Targeted Intervention, UCL Division of Surgery and Interventional Science, Charles Bell House, London, UK; Mitochondrial Research Group, Genetics and Genomic Medicine, UCL Great Ormond Street Institute of Child Health, and Metabolic Unit, Great Ormond Street Hospital for Children NHS Foundation Trust, London, UK

## Abstract

Pyruvate dehydrogenase complex deficiency (PDHD) is a severe mitochondrial disorder most frequently caused by pathogenic variants in *PDHA1,* leading to neurodevelopmental delay and early mortality, necessitating brain-targeted interventions. Using a brain-specific *Pdha1* knockout mouse model, we compared intracerebroventricular delivery of AAV9 capsid and a recently described synthetic neurotropic AAV-F capsid, both expressing human *PDHA1* coding sequence driven by a constitutive CAG promoter. Newborn mice received, titre matched AAV9 or AAV-F or AAV9 at ten-fold higher dose. Low-dose AAV-F and high-dose AAV9 significantly improved survival, and restored PDH enzyme activity, metabolite profiles, and brain histopathology to near wild-type levels. However, treated mice showed reduced locomotion by P100 and impaired motor function. Importantly, AAV-F achieved broad CNS transduction with minimal liver expression, outperforming AAV9 at lower dose. There results support the therapeutic potential of AAV-based gene therapy for PDHD and highlighting AAV-F as a promising capsid for efficient, CNS specific delivery.

## Introduction

Pyruvate dehydrogenase complex (PDHc) deficiency is a rare disorder of energy metabolism^1^. The PDHc enzyme complex converts pyruvate to acetyl-CoA, fuelling the Krebs cycle and the subsequent generation of ATP via oxidative phosphorylation in the mitochondria^2^. A deficiency of PDHc leads to accumulation of pyruvate and lactate, decreased ATP production^3^ and disrupted neurotransmitter synthesis, particularly acetylcholine, glutamate, and γ-aminobutyric acid (GABA)^4^. The resulting energy deficit particularly affects organs with high energy demands, such as the brain and muscles^5^, altering neuronal excitability and causing neurological symptoms^6^.

PDHc Deficiency (PDHD) is inherited in an X-linked or autosomal recessive manner, with most cases caused by pathogenic variants in the X-chromosomal *PDHA1* gene, which encodes the E1-alpha subunit of PDHc^7^. Symptoms vary based on the severity of the specific genetic variant, residual enzyme activity^8^, and patient’s sex. Hallmark features include central nervous system (CNS) abnormalities, developmental delay, hypotonia, ataxia, seizures^4^, encephalopathy, motor dysfunction, and severe lactic acidosis^9^. Neuroimaging often reveals ventriculomegaly and brain malformations such as hypogenesis of the corpus callosum^10^. Currently, a ketogenic diet is the recommended treatment and has been shown to improve motor function and reduce neuro-inflammation in patients with PDHD, but disease progression can continue^11,12^.

Adeno-associated virus serotype 9 (AAV9) based gene therapies are being explored and applied in numerous preclinical and clinical trials for genetic disorders, particularly those affecting the CNS and neuromuscular system^13–18^. A key advantage of AAV9 is its ability to cross the blood-nervous system barrier, and its ability to deliver to the nervous system and visceral organs makes it useful for treating multi-system diseases such as neurometabolic diseases^13^. With high transduction of motor neurons, AAV9 has also shown great clinical efficacy in Zolgensma for spinal muscular atrophy^19^. High doses and volumes of AAV9 are required to achieve the desired therapeutic effects, but such high doses have been associated with off-target toxicities and immune responses^20,21^.

Synthetic modifications to naturally occurring capsids have allowed the development of novel capsids with greater transduction efficacy^22^. By the introduction of a ligand or peptide at a specific site on the capsid surface, potency and tropism are improved, leading to efficient targeting. These improvements can achieve a reduction in immunogenicity, as doses can be lowered without affecting the therapeutic effect. AAV-F, a novel synthetic AAV capsid derived from AAV9, has been constructed with the aims of increasing tropism to the CNS and reducing the toxicity and immune responses associated with the parental capsid. AAV-F was generated by introducing a 7-amino acid insertion into the AAV9 Cap sequence. AAV-F demonstrated a greater CNS transduction profile after tail vein injections to adult mice^23^. When translated to non-human primates (NHP), AAV-F had a greater vector copy number (VCN) in spinal cord than titre matched AAV9 after intrathecal delivery^24^. In addition, AAV-F has recently been used in a pre-clinical study to treat Dravet syndrome^25^.

In this study we used a brain specific knock-out (KO) mouse model for PDHD, which closely recapitulates the human disease. The model was generated by deletion of exon 8 of the X chromosomal *Pdha1* gene^9^ and previous studies have demonstrated stunted growth, epilepsy, neuropathology (neuronal loss, increased GFAP, cortical thinning and hypoplasia of the corpus callosum), reduction of Pdha1 in neurons and astrocytes, and early death, specifically in male *Pdha1* KO mice^5,26^. In addition, we have shown that the model presents abnormal locomotion and motor defects. We aimed to develop a one-off AAV-F mediated gene supplementation therapy to deliver human *PDHA1* gene to the neonatal male *Pdha1* KO mice via unilateral intracerebroventricular (ICV) administration. We demonstrated that AAV-F gene therapy improved weight, survival, behavioural assessments, enzyme activity, metabolites and neuropathology in treated *Pdha1* KO mice.

## Results

### Widespread CNS gene expression after neonatal intracerebroventricular administration of AAV-F to wild-type mice

To assess the expression profile of AAV-F after neonatal delivery, we administered AAV-F-CAG-eGFP at low dose (LD; 5×10^9^ vg/pup) and AAV9-CAG-eGFP at LD and high dose (HD; 5×10^10^ vg/pup) to newborn wild-type (WT) mice via unilateral intracerebroventricular (ICV) injections (Fig.1A). Five weeks later, we directly observed extensive green fluorescence protein (GFP) expression in the brain and visceral organs of AAV-F and AAV9 treated groups (Supplementary Fig. 1A). Immunohistochemistry revealed comparable GFP expression throughout the brain with AAV-F-CAG-eGFP LD and AAV9-CAG-eGFP HD (Supplementary Fig. 1B). Quantification of immunoreactivity of GFP showed that AAV-F LD and AAV9 HD had comparable GFP expression in the prefrontal cortex, cortex, striatum, hippocampus, midbrain and cerebellum (Supplementary Fig. 1C).

**Fig. 1.**
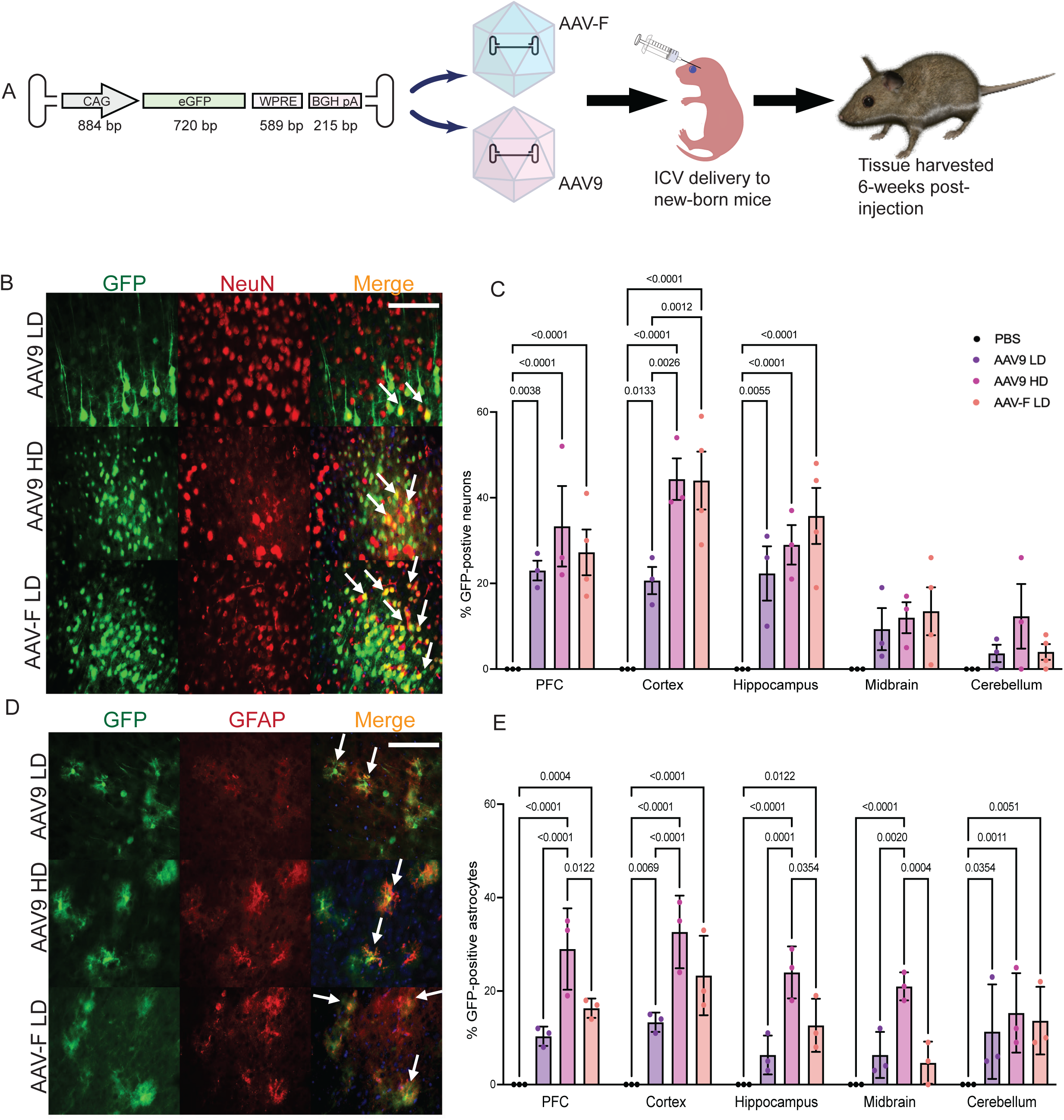
AAV-F mediates a greater transduction efficiency of neurons and astrocytes than titre-matched AAV9 after neonatal ICV delivery. **(A)** Schematic diagram of neonatal biodistribution study. **(B)** Representative images of co-localisation of NeuN and GFP in the cortex. Scale bar shown (100µm). **(C)** Stereological quantification of NeuN and GFP Immunofluorescence co-stain. One-way ANOVA, Tukey’s multiple comparison, with ± SEM (n=4). **(D)** Representative images of co-localisation of GFAP and GFP in the cortex. Scale bar shown (100µm). **(E)** Stereological quantification of GFAP and GFP Immunofluorescence co-stain. One-way ANOVA, Tukey’s multiple comparison, with ± SEM (n=4).

Co-localisation studies were performed to determine neuronal and astrocyte targeting of AAV9 and AAV-F capsids in the brain after neonatal ICV administration. AAV-F LD treated mice demonstrated comparable GFP-positive neurons in the prefrontal cortex (27.25%), cortex (44%), and hippocampus (35.7%) with AAV9 HD (33.3%, 44.3%, 29%; respectively) (Fig. 1B & C). AAV-F LD showed a significantly higher percentage GFP-positive neurons in the cortex compared to titre matched AAV9 (p=0.0012) (Fig. 1C). Astrocyte targeting was also observed with both AAV-F and AAV9 vectors (Fig. 1D). Notably, AAV9 HD showing a significant astrocytic transduction in the prefrontal cortex (29%), hippocampus (18%), and midbrain (21%), when compared to AAV9 LD (10.3%, 6.3%, 6.3%; p<0.0001) and AAV-F LD (16.3%; p=0.0122, 12.6%; p=0.0354, 4.6%; p=0.0004) (Fig.1E).

### AAV-F-CAG-*hPDHA1* vector targets the brain with an acceptable safety profile

To assess the biodistribution and safety of AAV-F-CAG-*hPDHA1* vector (Fig. 2A) we administered AAV-F-CAG-*hPDHA1* LD and AAV9-CAG-*hPDHA1* LD and HD vectors to newborn WT mice via unilateral ICV injections. We observed no difference in weight of treated groups compared to controls (Supplementary Fig. 2). Six weeks post-injection, VCN, *PDHA1* mRNA expression in the brain and visceral organs and neuropathology were assessed.

**Fig. 2.**
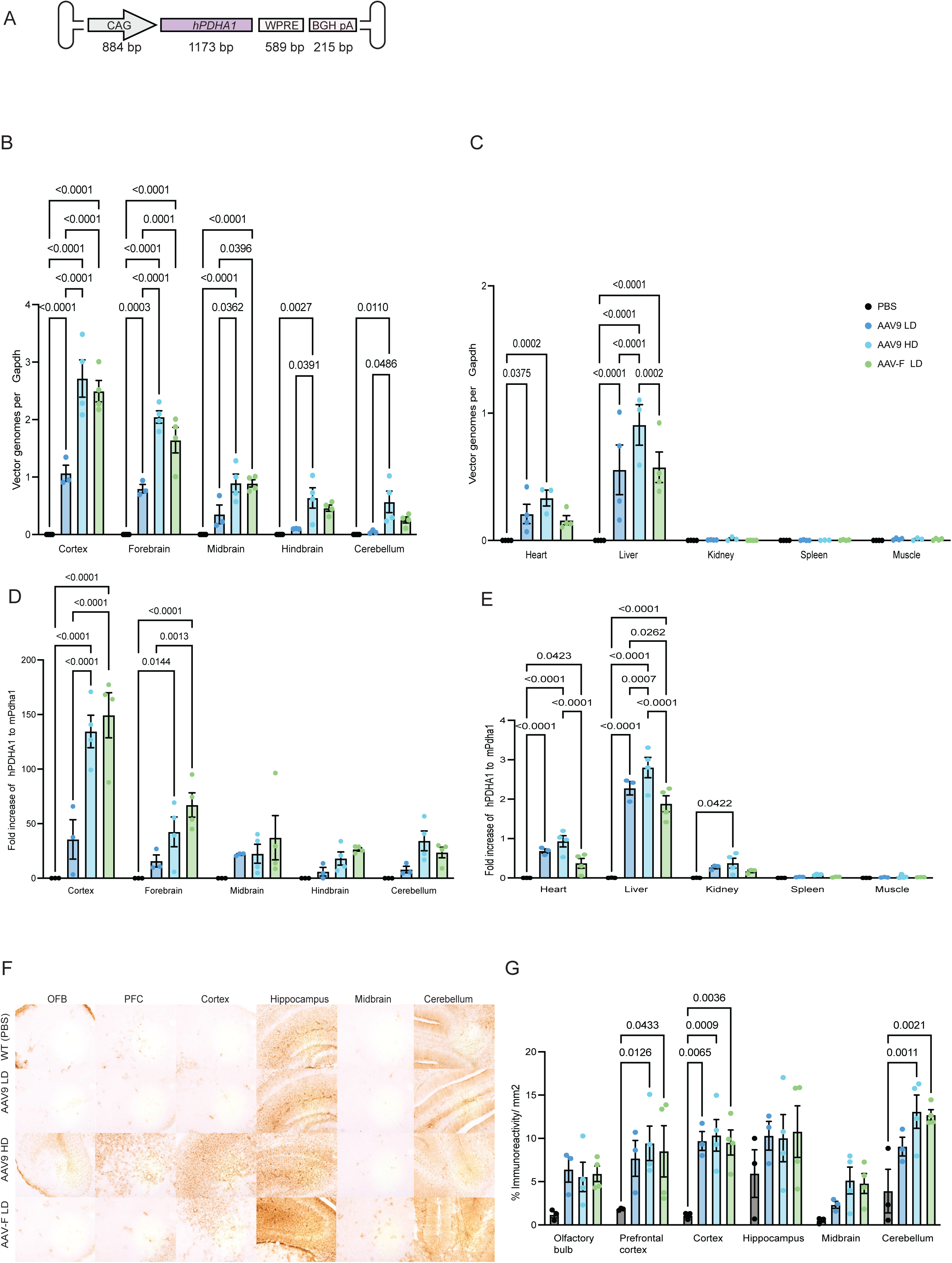
Neonatal intravenous delivery of AAV-F extensively transduces the CNS and visceral organs in WT mice. **(A)** Schematic diagram of therapeutic plasmid containing *hPDHA1.* **(B)** Vector genome analysis in discrete regions of the brain. One-way ANOVA, Tukey’s multiple comparison, ± SEM (n=4). **(C)** Fold increase of *hPDHA1* to *mPdha1* (normalised to *mGapdh)* in discrete regions of the brain. One-way ANOVA, Tukey’s multiple comparison ± SEM (n=4). **(D)** Vector genome analysis of visceral organs. One-way ANOVA, Tukey’s multiple comparison with ± SEM (n=4). **(E)** Fold increase of *hPDHA1* to *mPdha1* (normalised to *mGapdh)* in visceral organs One-way ANOVA, Tukey’s multiple comparison, ± SEM (n=4). **(F)** Representative images of GFAP expression in WT mice OFB, PFC, cortex, hippocampus, midbrain and cerebellum. One-way ANOVA, with Dunnett’s multiple comparison ± SEM. Images were taken at 400X magnification. Scale bar = 100µm. **(G)** Quantification of GFAP expression in the olfactory bulb (OFB), prefrontal cortex (PFC), cortex, striatum, hippocampus, midbrain and cerebellum ± SEM (n=4).

VCN assessment was comparable for AAV9 HD and AAV-F LD groups throughout the brain (Fig. 2B). AAV-F LD revealed a significant increase in VCN in the cortex, forebrain and midbrain compared to titre matched AAV9 LD. VCN analysis in the visceral organs showed significantly greater VCN in the liver of AAV9 (0.9) HD group compared to titre matched AAV-F LD (0.57, p=0.0002) and AAV9 LD (0.55, p<0.0001) (Fig. 2C). Comparable VCN were observed in the heart for all vectors and doses.

*PDHA1* expression following AAV-F LD showed a significant increase in the cortex (p<0.0001) and the forebrain (p=0.0004) only when compared to titre matched AAV9 (Fig. 2D). There was no significant difference throughout the brain between AAV-F LD and AAV9 HD (Fig. 2D). In the visceral organs, AAV-F LD displayed no significant difference in *PDHA1* expression compared to titre matched AAV9 (Fig. 2E). AAV9 HD had significantly higher *PDHA1* expression compared to AAV9 LD and AAV-F LD groups in the liver (AAV9 LD; p=0.0007, AAV-F LD; p<0.0001) (Fig. 2E).

Immunohistochemistry for glial fibrillary acidic protein (GFAP; marker for astrocytes) and CD68 (marker for active microglia) demonstrated significant increase in GFAP expression in the pre-frontal cortex (AAV9; p=0.0126, AAV-F; p=0.0433), cortex (AAV9; p=0.0009, AAV-F; p=0.0036) and cerebellum (AAV9; p=0.0011, AAV-F; p=0.0021) by AAV9 HD and AAV-F LD compared to PBS control group (Fig. 2F & G). GFAP expression was significantly increased in AAV9 LD group in the cortex only (p=0.0065) compared to PBS control group.

### AAV-F-CAG-*hPDHA1* improved survival and neurological phenotype of *Pdha1* KO mice

Prior to testing the efficacy of our novel gene therapy, we first aimed to further characterise the phenotype of the *Pdha1* KO mouse model (Supplementary Fig. 3A-C). For the first time, we observed that this model presents with abnormal locomotion as measured by behavioural tests, rotarod and open field activity (Supplementary Fig. 3D-G). There was a significant reduction in *mPdha1* expression, compared to age-matched WT controls (Supplementary Fig. 3H). Previous neuropathology assessment reported co-localised Pdha1 expression with GFAP and neurons (in the cortex, hippocampus and cerebellum), cortical thinning and hypoplasia of the corpus callosum^5,26^. Here we observed widespread over expression of GFAP and neuronal loss in the brain (Supplementary Fig. 3I & J).

Next, we sought to assess the efficacy of AAV-F-CAG-*hPDHA1* gene therapy treatment in male *Pdha1* KO mice. AAV-F-CAG-*hPDHA1* LD was administered via unilateral ICV injections, using AAV9-CAG-*hPDHA1* LD and HD as comparators. PBS was delivered via unilateral ICV injections to *Pdha1* KO mice and WTs as control groups. We monitored the mice for 100 days of development. We observed 80% survival for AAV9 HD (p=0.14), 75% for AAV-F LD (p=0.1), 60% for AAV9 LD (p=0.02), and 0% for KO PBS mice (p<0.001) compared to 100% survival of WT PBS mice (Fig. 3B). There was a significant difference in weight between all treatment groups compared to the control PBS KO group (p=0.0005) (Fig. 3C). There was no significant difference in weight between the PBS WT group and mice that received both AAV9 LD (p=0.9409), AAV9 HD (p=0.9235) and AAV-F LD (p=0.9473) gene therapy at the end of the in-life experiment (Fig. 3C).

**Fig. 3.**
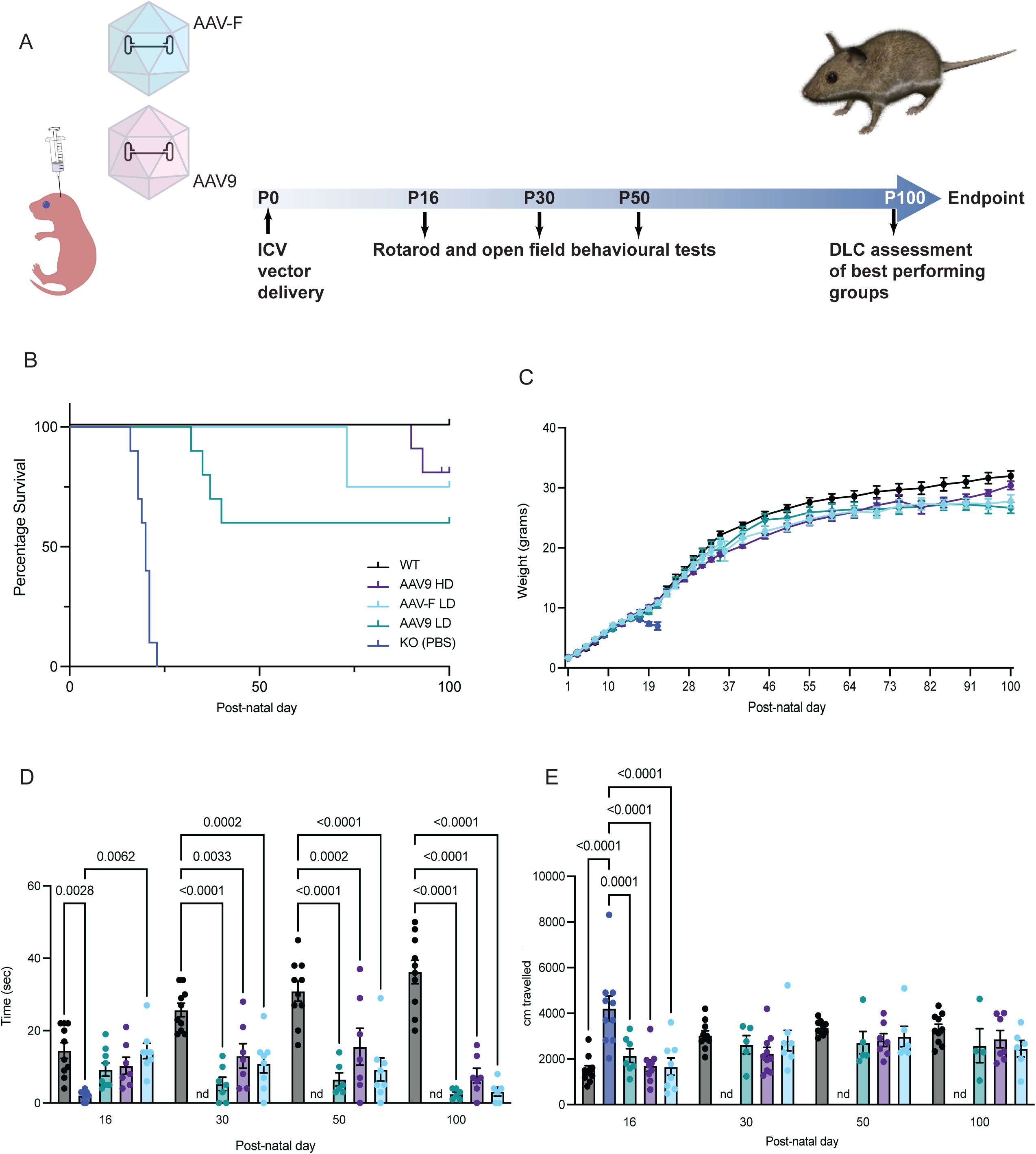
AAV-F gene therapy improves disease symptoms in a PDHD mouse model. **(A)** Schematic diagram illustrating the pre-clinical gene therapy study, **(B)** Kaplan-Meier survival curve. Data presented as a percentage of survival (n=8 for AAV-F LD, n=10 for all other groups) **(C)** Weight shown as mean ± SEM. **(D)** Behavioural rotarod assessment, showing time spent on accelerated rotarod. One-way ANOVA, Tukey’s multiple comparison ± SEM. **(E)** Open field test showing total distance travelled in 15 minutes. One-way ANOVA, Tukey’s multiple comparison ± SEM. Not done (nd).

Rotarod and open field behavioural assessments were performed at P16, P30, P50 and P100 to assess improvements in motor coordination and locomotion after AAV-F and AAV9 gene therapy. The rotarod test at P16 demonstrated that only the AAV-F LD (P=0.0062) and PBS WT (p=0.0028) groups had a significant improvement in time spent on the rotarod compared to the PBS KO group (Fig. 3D). As the PBS KO mice died at P21/22 of development, the remaining behavioural time points were compared to PBS WT mice. Rotarod at P30, P50 and P100 revealed no significant improvement in motor coordination in all AAV-F and AAV9 treatment groups compared to PBS treated WT mice (Fig. 3D). In contrast, open field assessment revealed a significant improvement at all doses for AAV-F and AAV9 at P16 compared to PBS treated KO mice (Fig. 3E). Open field analysis at remaining time points showed comparable results between all AAV-F and AAV9 groups and the PBS WT control group (Fig. 3E).

As open field assessment failed to detect nuanced behavioural and motor abnormalities, novel AI assessment using DeepLabCut (DLC) was applied. DLC was used to obtain high-resolution, marker-less tracking of specific body parts, enabling the identification of fine-grained phenotypic features such as gait irregularities, altered posture, or impaired limb coordination. This approach allowed for a more sensitive and quantitative assessment of both disease-related motor deficits and treatment-induced improvements that were not apparent through conventional open field metrics (Fig. 4A-D).

**Fig. 4.**
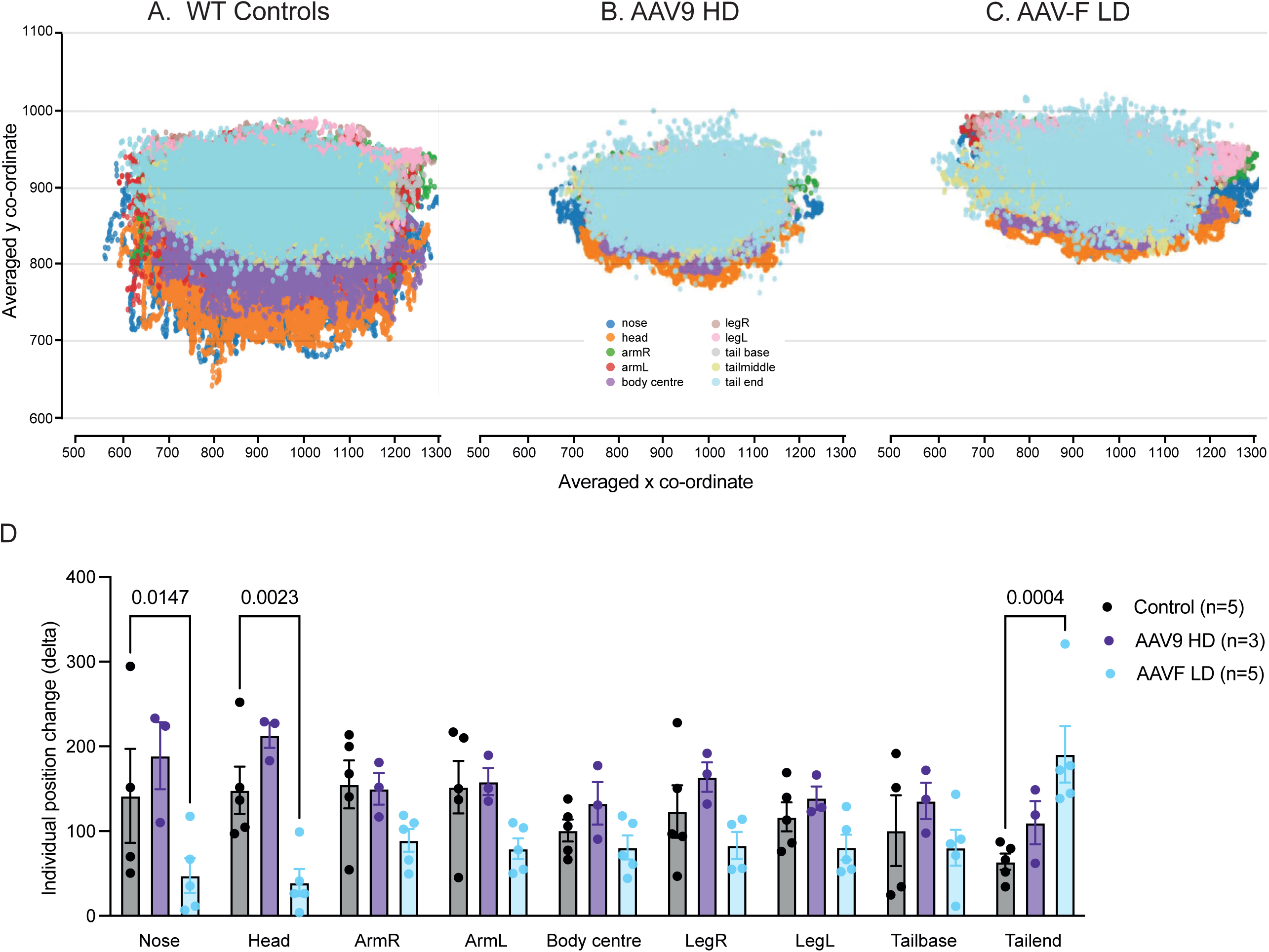
AI-behavioural assessment identifies abnormal phenotype. **(A)** Trajectory plot showing behavioural pose estimation of PBS-treated WT mice at postnatal day 100 (P100), recorded over a 15-minute video period. Each body part is represented by a unique colour: nose (blue), head (orange), right arm (green), left arm (red), body centre (purple), right leg (dark purple), left leg (light pink), tail base (grey), tail middle (light green), and tail end (light blue); This colour coding applies consistently across all trajectory graphs. **(B)** Trajectory plot for AAV9-HD-treated KO mice. **(C)** Trajectory plots showing AAV-F LD treated KO mice. **(D)** Quantification of Body-Part-Specific Movement Based on X-Y Coordinates from Figure 4B. One-way ANOVA, Dunnett’s multiple comparison.

This assessment was performed on mice treated with the best performing vector and dose, AAV-F LD and AAV9 HD at P100. WT control mice showed more spread and variability, possibly suggesting more free movement (Fig. 4A). The scatter plot for AAV9 HD and AAV-F LD revealed a tighter cluster of distribution with constrained movement of head and tail. This indicated that the treated mice continued to exhibit an abnormal phenotype relative to age-matched WT controls (Fig. 4B & C).

To quantify movement, we developed an algorithm calculating total displacement based on x and y coordinates in Fig. 4A-C (Supplementary Fig. 4). Figure 4D shows that while no significant differences were found for nose and head movements between the control and AAV9-HD groups, the AAV-F-LD group showed significantly reduced movement. Tail displacement was notably higher in the AAV-F-LD group than in the other groups (Supplementary Video 1).

Long-term improvements in PDHc enzyme activity and metabolites in the brain following gene therapy.

To assess the biodistribution of the vector, we examined *PDHA1* expression in discrete regions of the brain and visceral organs for all treatment groups. This was calculated based on fold-change of human *PDHA1* to mouse *Pdha1* (Supplementary Fig. 5A & B). We observed a significant increase in *PDHA1* expression in the cortex and forebrain following AAV9 HD and AAV-F LD (MD p<0.0001) (Fig. 5A). *PDHA1* expression was significantly higher in the liver AAV9 HD compared to AAV-F LD (p=0.0052). The heart revealed significant targeting with all vectors and doses (Fig. 5B).

**Fig. 5.**
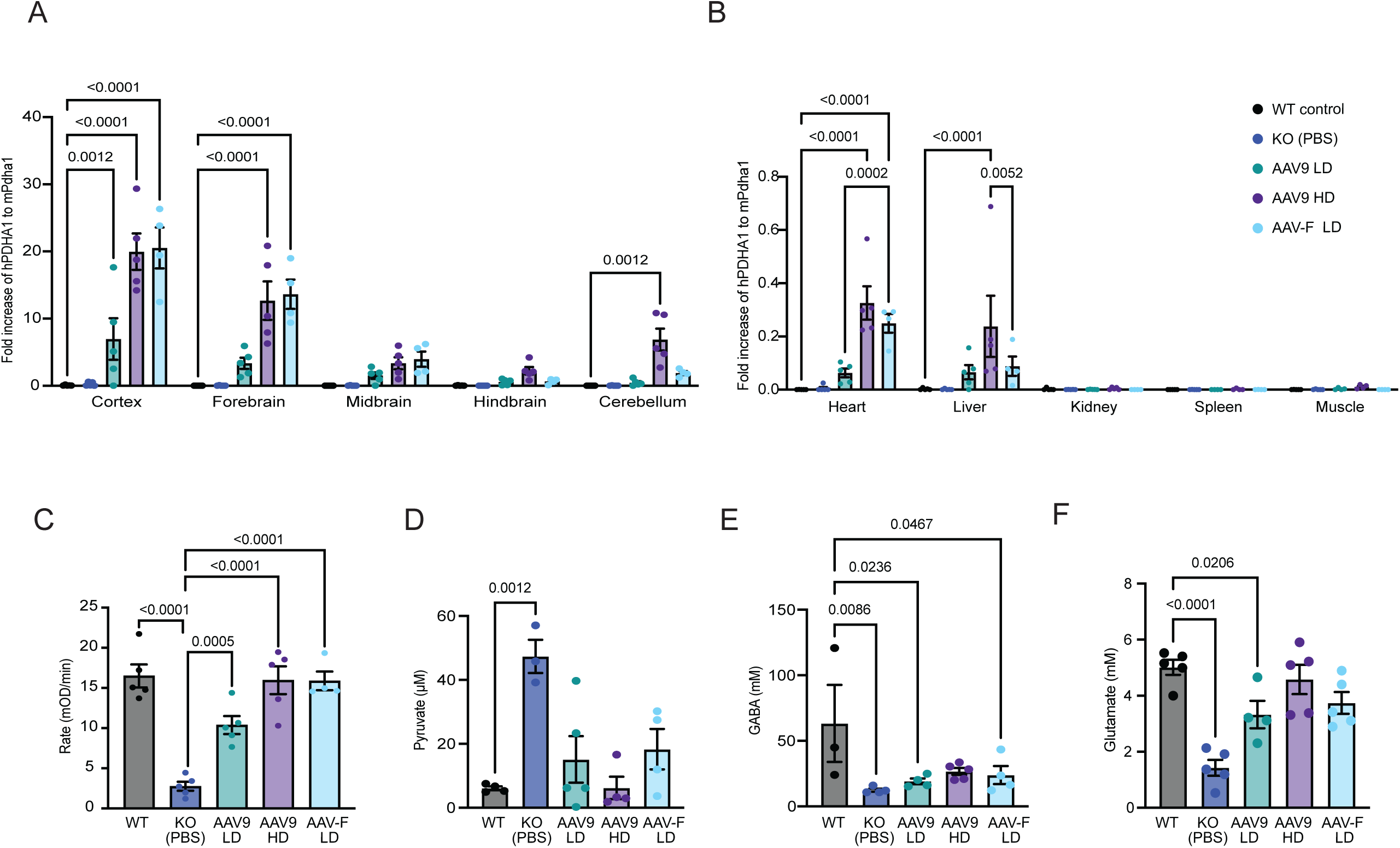
AAV-F LD is significantly more efficient at restoring biochemical phenotype to WT levels than titre-matched AAV9. **(A)** Fold increase of *hPDHA1* to *mPdha1* (normalised to *mGapdh)* in discrete brain regions. One-way ANOVA, Tukey’s multiple comparison ± SEM (n=5). **(B)** Fold increase of *hPDHA1* to *mPdha1* (normalised to *mGapdh)* in visceral organs. One-way ANOVA, Tukey’s multiple comparison ± SEM (n=5). **(C)** PDHc enzyme activity in in discrete regions of the brain, One-way ANOVA, Tukey’s multiple comparison ± SEM (n=5). **(D)** Pyruvate concentration in the brain. One-way ANOVA, Tukey’s multiple comparison ± SEM (n=5). **(E)** Glutamate concentration in the brain. One-way ANOVA, Tukey’s multiple comparison ± SEM (n=5). **(F)** GABA concentration in the brain. One-way ANOVA, Tukey’s multiple comparison ± SEM (n=5).

We observed significantly increased PDHc enzyme activity in AAV9 HD (15.95 mOD/min; p<0.0001), AAV-F LD (15.88 mOD/min; p<0.0001), and AAV9 LD (10.37 mOD/min; p=0.0005), compared to KO PBS treated mice (2.5 mOD/min). AAV9 HD and AAV-F LD were comparable to PBS WT controls (16.29 moD/min) (Fig. 5C).

We also assessed pyruvate concentrations in the brains of treated *Pdha1* KO mice as a biomarker for PDHD. Pyruvate levels were rescued to WT levels (6.1 µM) by AAV9 LD (15.1 µM), AAV9 HD (6.2 µM) and AAV-F LD (18.3 µM), with KO PBS treated mice displaying 45.8 µM (Fig. 5D).

Levels of the neurotransmitters glutamate and GABA were quantified, since their concentrations are commonly altered as a result of PDH deficiency^4^. Glutamate levels were normalised in AAV9 HD (5482.64 µM) and AAV-F LD (3742.54 µM) which were not significantly different to PBS WT (5015.02 µM) or PBS KO (1427.8 µM) (Fig. 5E). AAV9 LD (p=0.026) showed a significantly lower concentration of glutamate compared to WT controls. GABA was restored to PBS WT levels (63297 µM) by AAV9 HD (26678 µM) only (Fig. 5F).

### AAV-F mediated gene therapy ameliorates neuropathology throughout the brain except for the cerebellum

To determine whether our novel AAV-F mediated gene therapy ameliorated the neuropathology in the *Pdha1* KO mice, we conducted GFAP, CD68, NeuN (marker for neuronal loss) and H&E immunohistochemistry on brain tissue from AAV-F and AAV9 treated mice.

The results showed that there was a reduction in the upregulation of GFAP in AAV-F LD, AAV9 LD and AAV9 HD throughout the brain except for the cerebellum, where titre matched AAV-F LD (p=0.0011) and AAV9 LD (p<0.0001) displayed a significant elevation of GFAP positive cells compared to WT controls (Fig. 6A & B).

**Fig. 6.**
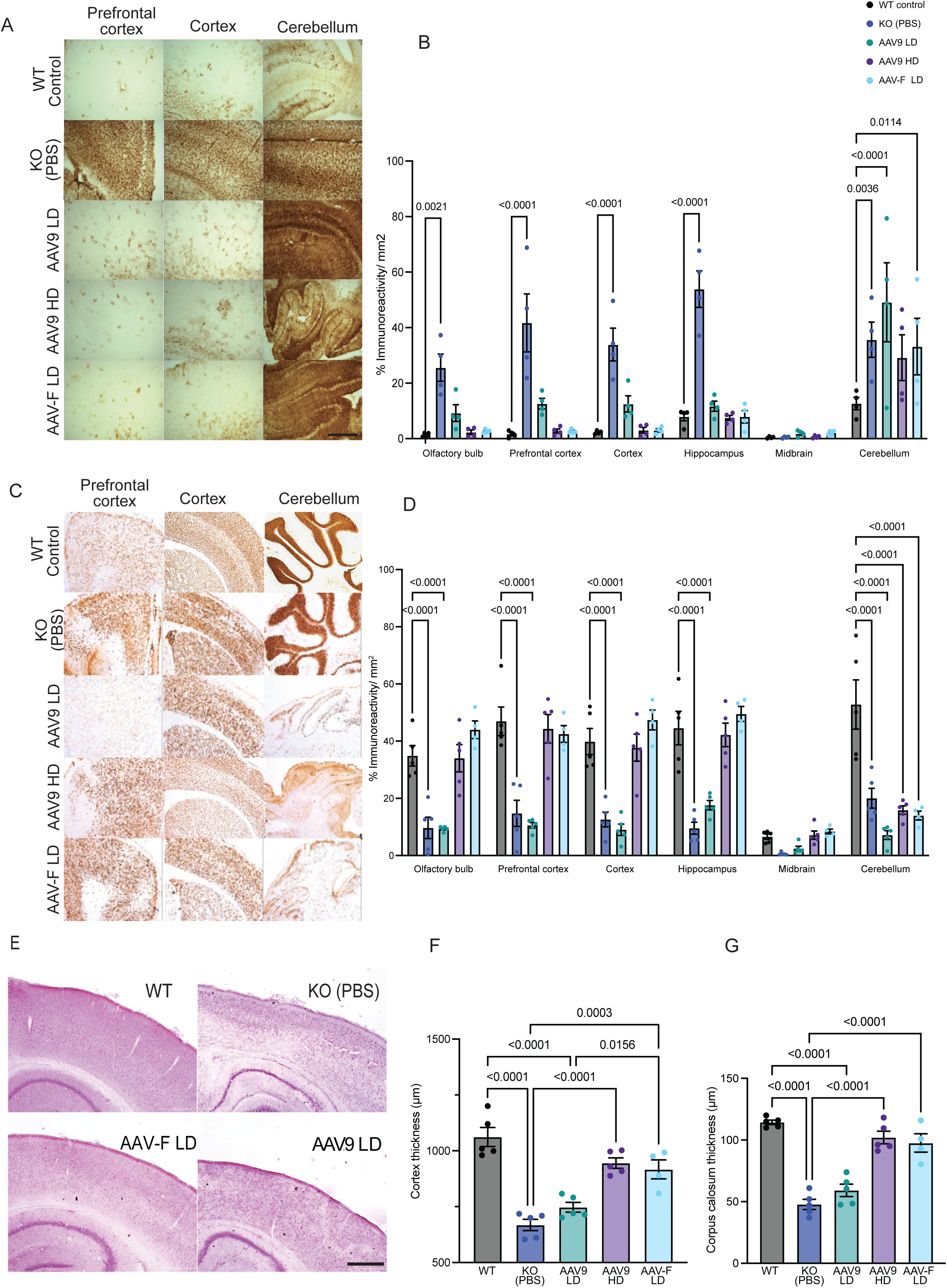
AAV-F LD improves neuropathology in PDHc deficient mice compared to titre-matched AAV9. **(A)** Representative images of GFAP expression in the prefrontal cortex (PFC), cortex and cerebellum. **(B)** Quantification of GFAP expression in the olfactory bulb (OFB), prefrontal cortex (PFC), cortex, hippocampus, midbrain and cerebellum. One-way ANOVA, Tukey’s multiple comparison ± SEM. **(C)** Representative images of neuronal loss in the prefrontal cortex (PFC), cortex and cerebellum. **(D)** Quantification of neuronal loss in the olfactory bulb (OFB), prefrontal cortex (PFC), cortex, hippocampus, midbrain and cerebellum. One-way ANOVA, Tukey’s multiple comparison ± SEM. **(E)** Representative images of H&E staining of the cortex. **(F)** Quantification of cortex thickness. One-way ANOVA, Tukey’s multiple comparison ± SEM. **(G)** Quantification of corpus callosum thickness. One-way ANOVA, Tukey’s multiple comparison ± SEM.

NeuN staining demonstrated a comparable neuronal immunoreactivity in the olfactory bulb, pre-frontal cortex, cortex and hippocampus in AAV-F LD and AAV9 HD compared to PBS WT group. However, neuronal immunoreactivity was significantly reduced in the cerebellum for both AAV-F LD (p<0.001) and AAV9 HD (p<0.001) compared to PBS WT controls (Fig. 6C & D). AAV9 LD showed significant reduction in neurons in the olfactory bulb, pre-frontal cortex, cortex, hippocampus and cerebellum (p<0.0001) (Fig. 6D). CD68 stain showed no microglia activation in all treatment groups, throughout the brain (Supplementary Fig. 5C).

H&E stain was performed to measure cortical and corpus callosum thickness, as thinning of these are two biomarkers of neuropathology in both the *Pdha1* KO mice and patients with PDHD^10^. Cortical thinning was improved for both AAV-F LD and AAV9 HD (Fig. 6E). Cortical thinning was significantly improved for both AAV-F LD (p=0.003) and AAV9 HD (p<0.001) compared to PBS-treated KO controls (Fig. 6F). Cortical atrophy was not restored by AAV9 LD as there was no significant difference from PBS-treated KOs. In the corpus callosum, thinning was restored to WT levels by AAV-F LD and AAV9 HD treatment, whereas AAV9 LD-treated mice exhibited thinning similar to KO controls (Fig. 6G). We demonstrated a significant improvement in cortical and corpus callosum thickening in AAV-F LD and AAV9 HD treated animals compared to PBS KO mice. Our findings suggest that our new therapy facilitates anatomical restoration and gives protection against neurodegeneration in PDHc deficient mice.

## Discussion

This study demonstrated for the first time that our proof-of-concept AAV-F-mediated gene therapy can ameliorate the severe disease phenotype in a clinically relevant *Pdha1* KO mouse model, highlighting the potential of a one-off AAV-F gene therapy for PDHD.

AAV vectors, specifically AAV9, have been used extensively to treat neurological disorders including spinal muscular atrophy^19^. However, high doses of AAV9 have been linked to liver toxicity and thrombotic microangiopathy^27^. Novel capsids that enable efficient therapeutic delivery at a lower dose are needed, to avoid immunotoxicity associated with gene therapy. A novel AAV-F capsid has shown greater CNS transduction efficiency than titre matched parental AAV9 capsid^23,24^.

In our proof-of-concept study we have shown that AAV-F gene therapy to the brain of newborn PDHc deficient mice at a lower dose than AAV9 (ten-fold higher dose than AAV-F) can significantly extend life, reduce neurological disease symptoms, and restore PDHc enzyme activity and metabolites in the brain.

Prior to administering the *PDHA1* therapeutic vector to WT or *Pdha1* KO mice, we established the expression profile of AAV-F compared to titre matched AAV9 expressing eGFP under the control of CAG promoter, after single unilateral ICV administration to WT neonatal mice. An additional dose of AAV9 at one log higher was also administered. This study revealed that the novel-engineered AAV-F capsid had enhanced neurotropism and CNS transduction rate compared to the parental AAV9 capsid, five weeks after a single neonatal ICV injection. Previous studies have shown a significantly greater transduction of neurons and astrocytes in the CNS after adult tail vein injection with AAV-F compared to titre matched AAV9^23^, thus aligning with our findings. Furthermore, the AAV-F capsid was engineered for CNS specificity, exhibiting enhanced brain tropism compared to its parental AAV9 capsid, likely attributable to improved transduction efficiency via the Ly6a receptor on murine brain endothelial cells ^23^. The Ly6a receptor was identified as a key target for novel AAV capsid binding^28^. The 7-mer peptide substitution in AAV-F enhances its binding affinity to Ly6a, thereby improving transduction efficiency^29^. Although Ly6a receptor is absent in human or primate brain endothelial cells, NHP studies have shown a higher VCN of AAV-F in the spinal cord compared to titre matched AAV9 after intrathecal delivery, thus indicating the potential for therapeutic use in larger animal species^24^.

We then administered AAV9 and AAV-F vectors expressing human *PDHA1* under the control of CAG promoter, to WT and *Pdha1* KO mice by single unilateral ICV injection. The rationale behind the use of ICV injection was based on treating the brain as the main priority due to the severe neuropathology identified in previous studies^26^. In addition, the CNS is the primary affected area in patients and therefore ICV injection would be the most applicable for clinical translation^30,31^. When the therapeutic vectors containing *hPDHA1* were administered to WT mice we observed no deaths and no statistical difference in weights compared to WT controls by the experimental endpoint. The vectors successfully delivered the *PDHA1* gene to the brain, with AAV-F showing higher VCN and more efficient gene expression, particularly in the cortex and midbrain. Interestingly, AAV-F at lower doses achieved similar expression to AAV9 at a higher dose, highlighting its potential for reduced vector dosing. While AAV9 HD had higher VCN in the cerebellum, this dose resulted in over-expression in the liver which was significantly higher than untreated-WT controls. AAV9 has broader tropism due to its ability to cross the blood brain barrier (BBB), its interaction with N-linked sialic acid receptors, and its systemic distribution^32^. Moreover, as the BBB remains naïve during the neonatal period the AAV vector has increased systemic targeting as it enters the bloodstream efficiently^33^. Liver targeting can be problematic in AAV-gene therapy due to the risk of hepatotoxicity^34^. These data suggest that by delivering AAV-F at ten-fold lower dose than AAV9 we can de-target the liver whilst maintaining high expression in the brain.

The *Pdha1* KO mouse model used in this gene therapy study was previously described by Jakkamsetti et al, with the *Pdha1* KO mice exhibiting stunted growth, epilepsy, neuropathology and early death^5^. The *Pdha1* KO mice also showed a down-regulation of PDHc enzyme activity and metabolite abnormalities (including glutamate and GABA), that closely recapitulate the human disease^7,35^. We further assessed this model and noted behavioural defects and neuropathology throughout the brain including upregulation of astrocytes and neuronal loss.

In the gene therapy study, growth outcomes in treated mice were significantly improved compared to PBS-treated KO controls. AAV9 HD and AAV-F LD mice had the largest improvement in survival of 80% and 75% respectively. Although ten-fold different in dose, there was no significant difference between either group’s survival. AAV9 LD revealed a 60% survival which was significantly lower than titre-matched AAV-F LD. Behavioural assessment also demonstrated a dose-dependent improvement at P16. Rotarod test showed that AAV-F LD treated mice spent longer on the rod than PBS-treated KO at P16. Although all treated groups had significantly higher latency to fall compared to WT controls at P30, P50 and P100, AAV9 HD had the longest time spent on the rod at all time points post-weaning. Open field studies showed similar results at P16 as there was no significant difference between the treated KOs and WT controls in total distance and mean speed. AAV9 LD and HD showed no significant decline in any of the open field assessments at P30, P50 or P100. AAV-F LD displayed a significantly lower mean speed at P100 compared to WT PBS controls. As the treated KO mice displayed an abnormal gait, which was identified in the rota rod assessment but undetected in the open field test, we aimed to identify another method which was more sensitive to assess the post treatment phenotype. Therefore, An AI-based behavioural analysis tool (DLC) was used to assess whether the two most effective treatment groups (AAV-F LD and AAV9 HD) showed phenotypic improvements, as well as to identify additional behavioural changes not detected by standard open field testing. DLC analysis revealed that AAV-F treated *Pdha1* KO mice continued to exhibit abnormal locomotor phenotypes compared to PBS-treated WT controls. Specifically, there was a significant reduction in nose, head, and tail-end movement in the AAV-F treated group. Impaired tail dynamics, as measured by DLC, have previously been used to assess the severity of ataxia in mouse models of spinocerebellar ataxia type 6 (SCA6)^36^. Tail movements are critical for balance in rodents, with active tail adjustments contributing significantly to postural stability, underscoring the tail’s key role in locomotion^37^. Likewise, deficits in balance and gait ataxia are common features in various neurodegenerative disease models^38,39^. Additionally, abnormal head and nose movements have also been identified in prior DLC studies, supporting the sensitivity of this tool in detecting subtle motor impairments^40,41^. Future studies are required to compare AAV9 HD and AAV-F HD and assess whether the increasing the dose of AAV-F can rectify the motor dysfunction observed in AAV-F LD group.

Brains of treated animals were assessed for gene expression, PDHc enzyme activity, Krebs cycle intermediates (pyruvate, GABA and glutamate) and neuropathology. These metabolites were selected as they are known be disrupted in the *Pdha1* KO model^5,26^. The cortex and the forebrain had the highest *hPDHA1* expression with both doses of AAV9 and AAV-F LD vector, while AAV9 HD showed the greatest cerebellar targeting. Neither AAV vectors altered gene expression in the midbrain and hindbrain. As we used a brain-directed approach, lower visceral organ targeting was expected. Although AAV9 HD and AAV-F LD displayed similar results in the cortex and forebrain, *hPDHA1* expression in the liver was significantly higher only in the AAV9 HD treated group. Interestingly AAV9 HD and AAV-F LD had a similar transduction in the heart. As there was little expression from AAV-F LD in the liver, it was not expected for the heart to have a significant increase. However, as these were neonatal ICV injections, the immature BBB has an increased permeability which allows an increase of cerebral blood flow^42^. The lack of rescue in visceral organs was not a priority of this study as visceral organ pathology is not a major clinical feature in human patients with PDHD.

It was important to evaluate whether gene therapy could restore PDHc enzyme activity, pyruvate, alanine, glutamate, and GABA to WT levels^5,26^. We showed that AAV9 HD and AAV-F LD restored PDHc enzyme activity to that of WT PBS treated mice. AAV9 HD ameliorated the abnormalities in the selected metabolites as it decreased pyruvate and increased the neurotransmitters glutamate and GABA. AAV-F LD was also successful but not to the same extent, as GABA levels remained the same as the PBS KO group. Titre matched AAV9 LD mice showed improvements in pyruvate, but glutamate and GABA concentrations were significantly lower than PBS WT controls. AAV9 HD likely outperformed AAV-F LD in restoring neurotransmitter levels because it achieved higher and more widespread CNS transgene expression particularly in critical neuronal subtypes responsible for glutamate and GABA synthesis. Future PDHc enzyme activity and metabolite assessment should be completed on discrete regions of the brain to indicate whether these disruptions are localised to the regions that lack therapeutic targeting.

The neuropathological analysis of the treated *Pdha1* KO mice confirmed that the neurological symptoms could be successfully ameliorated by AAV9 HD and AAV-F LD, throughout the brain except for the cerebellum. Both groups reduced the upregulation of astrocytes, neuronal loss, cortical thinning and hypoplasia of the corpus callosum. The observed improvements in cortical and corpus callosum thickness are likely a result of increased *PDHA1* expression in the cortex.

One limitation of this study was the lack of cerebellum targeting in all treatment groups. This results in an upregulation of GFAP astrocytes and reduced neuronal density. Despite ICV injections bypassing the BBB, the cerebellum sits below the fourth ventricle, and while the fourth ventricle is directly connected to the ventricular system, the cerebellum is further removed compared to other brain regions^43,44^. In addition, the cerebellum has a relatively different BBB compared to other brain regions and is not as well vascularised as the cerebral cortex, thus resulting in a reduction in vector delivery^45^. This absence of cerebellum targeting by this novel gene therapy requires further investigation; however, several pre-clinical studies have shown scarcity of expression in the cerebellum following ICV administration of AAV^46,47^. As there was no rescue of upregulation of astrocytes and neuronal death, disease progression continued as the mice advanced in age.

Overall, this study provides compelling evidence that AAV-mediated gene supplementation therapy rescues key disease pathological features in a the PDHc deficient mouse model. AAV9 HD and AAV-F LD significantly improved lifespan, abnormal brain architecture and neurodegeneration. Therefore, providing evidence that AAV gene therapy has the potential to offer a one-off long-term treatment for PDHD patients.

## Materials and Methods

### AAV production

AAV plasmid containing chicken beta-actin (CAG) promoter driving eGFP was aquired from Addgene (plasmid # 37825). AAV plasmid encoding the human *PDHA1* gene driven by a CAG promoter was ordered from GENWIZ LTD (Germany).

The recombinant AAV vectors, AAV-F (Addgene plasmid # 166921) and AAV9 (University of Pennsylvania, USA) were generated using the triple plasmid transfection method described previously^48^. AKTA high-performance liquid chromatography (HPLC) with POROS CaptureSelect AAVX Affinity Resin (Thermo Fisher Scientific) virus purification was performed. Vectors were titred via Taqman quantitative polymerase chain reaction (qPCR) by QuantStudio (Thermo Fisher Scientific).

### Animal research

All animal studies were performed according to the Animal (Scientific Procedures) Act 1986 and approved by the UK Home Office project license and Ethical Review Board (AWERB). Animals were housed in a controlled environment with a 12-hour light/dark cycle with food and water provided ad libitum. The biodistribution study was performed using wild-type C57BL/6J mice. The PDHc deficient model that was used was previously described by Jakkamsetti *et al*^5^ by crossing a *Pdha1* flox female (B6.129P2-Pdha1tm1Ptl/J; Jackson Laboratory: 017443) with a *hGFAP*-Cre male (FVB-Tg(GFAP-cre)25Mes/J; 004600). Prior to this crossing the *Pdha1* flox colony was moved onto a 129SV/J background.

ICV injections were performed on P0-P1 mice using a 33-gauge Hamilton needle, using coordinates described^49^. The biodistribution study used 5µl of titre-matched AAV-F and AAV9 (5×10^9^ vector genomes/pup). A third group was injected with AAV9 at a dose a log higher (5×10^10^ vector genomes/pup). The PDHc deficient model was administered with 5µl of titre-matched AAV-F and AAV9 containing the therapeutic gene (5×10^9^ vector genomes/pup) (low dose, LD). The third group received a dose of 5×10^10^ vector genomes/pup (high-dose, HD).

### Animal behavioural analysis

Behavioural assessments were performed on postnatal days 16, 30, 50 and 100. Three days prior to performing Rotarod test, animals were trained to walk on the rotating rod. On the day of testing the rotarod (Panlan LE8200, Cornella) was set to accelerate from 4 revolutions per minute (rpm) to 20rpm. The trial ended when the animal fell off the rod. The time and speed at fall were noted and the trial was repeated two more times. Trials were re-run if the animal fell off the rotating rod within the first 5 seconds of the start time.

Harvard Panlab open field-testing equipment was used to assess the activity and general movements (Barcelona, Spain). A camera was suspended from the ceiling facing four 25cm x 24cm open field arenas. The animals were each placed in one of the arenas where they were allowed to move freely for 15 minutes. The movements were analysed using SMART v3 software (Harvard Panlab). Total distance travelled, time spent in periphery/ centre, total resting time, mean speed and mean speed without resting were measured. The room was completely silent during recordings, with moderate lighting.

### AI assessment

The mice were placed in a perspex box (30cm x 30cm x 30cm) and recorded for 15 minutes at P100 of development. The videos were then imported into the DLC program. The DLC program and the code were used as described^50^ and DLC GitHub source (https://github.com/DeepLabCut). Additionally, customised codes were used for body-specific graphs and total body-specific positional change (Δγ=γfinal−γinitial=(Δx,Δy) analyses.

### Animal tissue harvesting

At the experimental endpoint, animals were placed under isoflurane anaesthesia and perfused with phosphate-buffered saline (PBS, Sigma-Aldrich) via the left ventricle of the heart. The animal was then euthanised and relevant organs were collected. The left hemisphere of the brain was fixed following collection in 4% paraformaldehyde solution (Sigma-Aldrich) for 48 hours before being moved to 30% sucrose for 24 hours. Other harvested tissues were stored at −80°C until required.

### qPCR analysis

DNA from tissue was extracted following the manufacturers’ protocol for Qiagen DNeasy Blood & Tissue kit (Qiagen). Analysis of vector copy number (VCN) per cell was determined using real-time quantitative PCR (qPCR) with Luna probe qPCR Master Mix (New England Biolabs). Primers were designed to target *mGapdh,* and *hPDHA1* (Supplementary Fig. 2C).

RNA was extracted using PureLink RNA purification kit (Thermofisher) and converted to cDNA with the Applied Biosystems high-capacity reverse transcription kit (Thermofisher). Fold change in gene expression per cell was determined using real-time quantitative PCR with Luna probe qPCR Master Mix (New England Biolabs, UK). RNA was Primers were designed to target *mGapdh, mPdha1* and *hPDHA1* (Supplementary Fig. 2C).

### PDHc enzyme activity

PDHc enzyme activity in the brain was measured using the PDHc enzyme activity microplate assay kit from Abcam UK (ab109902). One half of a frozen brain hemisphere was homogenised in PBS to extract protein before determining sample protein concentration using the bicinchoninic acid assay (BCA). 25µl of detergent supplied in the kit to the 500µl of sample and placed on ice for 10 minutes to solubilise. Following the incubation the samples were centrifuged at 3,000 rpm for 10 minutes at 4 °C. The supernatant was collected and the samples were then diluted to 1µg/µl in 400µl using 1X buffer.

200µl of sample was loaded into a well in the microplate and the plate was covered and incubated at room temperature for 3 hours. Following the 3-hour incubation period the wells were emptied and 300µl of 1X stabiliser was added to each well. Absorbance of each sample was read at 450 nm every 20 seconds for 30 minutes. The enzyme activity was identified as:

Rate (mOD/min) = ((A2 – A1) / (T2 - T1))/ 0.001

### Gas chromatography-mass spectrometry

The tissue collected was placed into a 15ml conical tube where it received 5ml of 5% acetic acid in methanol. The brain was homogenised at 20,000 rpm for 3 minutes using a mechanical homogeniser (IKA Labortechnick) and returned to an ice bath.

The homogenates were centrifuged at 2,000 rpm for 30 minutes at 4°C. The supernatant was then transferred to a clean tube and processed immediately or returned to −80°C until required. Internal standards of glutamate, alpha-ketoglutarate, pyruvate, beta-hydroxybutyrate and GABA were prepared as described previously. 100mM solution of methoxylamine-hydrochloride (MOX) at pH 8-10 was prepared fresh daily by adding 0.0082g of MOX to 10ml of dH20. This solution was then diluted to 4mM by adding 40μl to 960μl of dH20. 10μl of the 4mM MOX solution with 200μl of sample and 10μl of 1M internal standards was added to a glass vial and the pH was adjusted with 10M NaOH to ensure a pH of 8-10. The vials were incubated for 3 hours at 50°C on a heat block. After the 3 hours, the samples were transferred to a MAXI dry Iyo Heto Vacuum (MechaTech Systems, Bristol, UK) where they were vacuum dried overnight. The next day 100μl of MSTFA (Trifluoro-N-methyl-N-(trimethylsilyl)-acetamide (plus 1% TMCS (2,2,2-, Chlorotrimethylsilane) (Sigma-Aldrich) was added to the vial which was then incubated at 70°C for 50 minutes. Following the incubation, the samples were allowed to cool before transferring the 100μL to a 1mL gas vial. The sample was then injected into the gas chromatography-mass spectrometry (GC/MS) instrument. The machine used for GC/MS was ISQ™ 7610 Single Quadrupole (TRACE 1600) GC-MS (ThermoFisher Scientific). 2μl sample was injected at 250°C and a split ratio of 1:10. Oven temperature started at 45°C and was increased to 200°C at 10°C per minute and then to 300°C at 30°C per minute. The MS was operated at selected ion monitoring mode, monitoring for m/z 174 (pyruvate), 177 ^13^C_3_ (pyruvate internal standard), 191 (lactate), 116 (alanine), 233 (beta-hydroxybutyrate), 237 ^13^C_4_ (beta-hydroxybutyrate internal standard), 174 (GABA), 176 (d6-GABA internal standard, 246 (glutamate), 254 ^13^C_5_ (glutamate internal standard). Standards of 500, 250, 100, 50 25, 10 and 0 μM were set up using the same process mentioned previously.

### Immunofluorescence staining

Coronal 40µm brain sections were cut with a microtome (Thermo Fisher Scientific). Sections were treated with 1% hydrogen peroxide for 30 minutes, then washed three times in 1x Tris-buffered saline (TBS). Non-specific binding sites were blocked with 15% goat serum (Vector Laboratories) solution in 0.3% Triton X-100 TBS (TBS-T) for 30 minutes. Immunolabeling was performed on floating sections using antibodies directed against GFP (Abcam; ab290), NeuN (Merek; MAB377) and GFAP (Proteintech; 16825-1-AP) in 10% goat serum in TBST at 4°C overnight.

After 3 washes of 1xTBS, the sections were incubated in darkness with fluorescent secondary antibodies for 2 hours; goat anti-chicken Alexa Fluor 488 (against GFP, Thermofisher scientific; A-11039), goat anti-mouse Alexa Fluor 568 (against NeuN, Thermofisher scientific; A-11004) and goat anti-chicken Alexa Fluor 568 (against GFAP, Thermofisher scientific; A-11041) in 10% goat serum in TBS-T at room temperature. Following 3 washes of 1xTBS, DAPI stain (4′,6-diamidino-2-phenylindole) (Thermofisher scientific) was added for 5 minutes, and the sections were mounted onto double-coated gelatinised slides. Once dried, a cover slip was added using Fluromount (Ebioscience). Image analysis was performed on a fluorescence microscope (Leica DFC7000 7) using LAS X Microscope Software (Leica Microsystems). The percentage GFP positive neurons and astrocytes were analysed using the CellProfiler Software (Broad Institute 2021). The images were overlayed using ImageJ.

### Immunoperoxide staining

Coronal 40µm brain sections were subjected to 1% hydrogen peroxide for 30 minutes, then washed three times in 1x TBS. Non-specific binding sites were blocked with 15% goat serum (Vector Laboratories) solution in 0.3% Triton X-100 TBS (TBS-T) for 30 minutes. Immunolabeling was performed on floating sections using antibodies directed against GFP (Abcam), 10% goat serum in TBST at 4°C overnight. This protocol has been described by Karda et el^48^

### Hematoxylin and Eosin staining

Coronal 40µm brain sections were subjected to H&E staining, brains were prepared and stained as per the protocol previously described Massaro et al^51^.

### Statistics

Statistical analysis was performed on GraphPad Prism version 9.3.1, GraphPad Software, San Diego, California, USA (www.graphpad.com). P values that were less than 0.05 were taken to be statistically significant. Survival analysis was completed using the Mantel-Cox test. Weights were taken every other day and analysed with repeated measures ANOVA. One-way ANOVA with Tukey’s multiple comparison tests was used for GFP-positive neurons and astrocytes, VCN, rotarod, open field, qPCR, PDHc enzyme activity and neuropathology assessments. Data are presented as mean and standard error of the mean (SEM).

## Supporting information

Supplementary Method 1. DLC code

DLC Video - Control

DLC video - AAV9 HD

DLC video - AAV-F LD

## Acknowledgments

LifeArc P2020-0008 and P2023-0011 (R.K., J.A.D., E.C, S.N.W.). Great Ormond Street Hospital Children’s Charity and Dravet Syndrome UK Charity V4720 and V4919 (R.K., J.A.D., S.N.W., and E.C.). Therapeutic Acceleration Support (TAS), UCL (R.K., A.K., and S.R). Medical Research Council Development Pathway Funding Scheme MR/Z505201/1 (R.K. and E.C). Freya Foundation Charity.

## Author contributions

R.K., S.R., A.K., J.R.C, designed the research. R.K. and A.K. drafted the manuscript. A.K., O.C., J.A.D., E.C., performed the experimental studies, A.K., and O.C., analysed data. R.K., S.R., S.N.W., S.E., and A.K. guided research. All authors agreed the final version of this manuscript.

---

**Supplementary Fig. 1 GFP Biodistribution of AAV9 and AAV-F. (A)** Representative images of ex vivo GFP after neonatal ICV injections of AAV-F and AAV9. *Ex-vivo* GFP analysis at 100 days post injection. Scale bars shown in each image (2mm). **(B)** Representative images of GFP protein expression in the brains of PBS-treated controls, AAV9 LD and HD and AAV-F LD. Images were taken at 400X magnification. Scale bar = 100µm. **(C)** Quantification of GFP expression in the prefrontal cortex (PFC), cortex, striatum, hippocampus, midbrain and cerebellum. One-way ANOVA, Dunnett’s multiple comparison (mean± SEM). **(D)** Transduction of neurons following the administration of AAV9 LD and HD and AAV-F LD. Co-immunofluorescence with neuronal marker NeuN, GFP and nuclear stain, DAPI. Representative images of prefrontal cortex (PFC), cortex, hippocampus, midbrain and cerebellum. Images were taken at 400X magnification. Scale bar = 100µm. **(E)** Transduction of astrocytes following the administration of AAV9 LD and HD and AAV-F LD. Co-immunofluorescence with astrocyte marker GFAP, GFP and nuclear stain, DAPI. Representative images of prefrontal cortex (PFC), cortex, hippocampus, midbrain and cerebellum. Images were taken at 400X magnification. Scale bar = 100µm.

**Supplementary Fig. 2 WT toxicology study of therapeutic vector following ICV delivery of AAV9 and AAV-F. (A)** Weights of WT treated mice shown as mean ± SEM. One-way ANOVA multiple comparison. **(B)** Table of primers, probes and sequences used in this study.

**Supplementary Fig. 3 Characterisation of untreated PDHc deficient mice. (A)** Kaplan-Meier survival curve of KO mice compared to healthy controls (n=10). **(B)** Weights of KO mice compared to healthy controls (n=10). **(C)** Image of PDHc deficient mouse (right) alongside WT litter mate (left). **(D)** Rotarod assessment plotted as time (seconds) spent on the accelerating rod. *Pdha1* KO mice had a significantly higher latency to fall compared to healthy age matched littermates. **(E)** Open field test showing mean speed over 15 minutes. Welch’s t-test ± SEM. **(F)** Open field test showing total distance travelled in 15 minutes. Welch’s t-test ± SEM. **(G)** Open field test quantifying the resting time in 15 minutes. Welch’s t-test ± SEM. **(H)** Fold change of *Pdha1* gene expression to *Gapdh* in *Pdha1* KO mice compared to WT littermates.GFAP expression in *Pdha1* KO brain compared to WT littermates showing an upregulation of astrocytes throughout the brain. **(I)** NeuN expression in *Pdha1* KO brain compared to WT littermates showing extensive neuronal loss.

**Supplementary Fig. 4. Individual body part assessment using DLC. (A)** Targeted body part separated trajectory plots representing the PBS-treated control group. In all graphs, each body part is consistently color-coded as follows: nose (blue), head (light blue), right arm (dark orange), left arm (light orange), body centre (dark green), right leg (light green), left leg (red), tail base (pink), and tail end (purple). **(B)** Targeted body part separated trajectory plots representing AAV9-HD-treated KO mice show a change in individual body part movements compared to control mice. **(C)** Targeted body part separated trajectory plots representing AAV-F-LD treated KO mice show a change in individual body part movements compared to control mice.

**Supplementary Fig. 5. Further assessment of PBS-treated WT controls, and PBS treated, AAV9-LD and HD and AAV-F LD treated KO’s. (A)** Endogenous *Pdha1* expression in discrete regions of the brains of the treated KO mice. One-way ANOVA, Tukey’s multiple comparison ± SEM. **(B)** Endogenous *Pdha1* expression in visceral organs of the treated KO mice. One-way ANOVA, Tukey’s multiple comparison ± SEM. **(C)** Representative images of CD68 expression in KO-treated mice OFB, PFC, cortex, hippocampus, midbrain and cerebellum. Images were taken at 400X magnification. Scale bar = 100µm.

**Supplementary Video 1. Videos of treated KO *Pdha1* mice labelled with body parts. (A)** Control mouse. **(B)** AAV9 HD treated mouse. **(C)** AAV-F LD treated mouse.

**Supplementary Method 1. DLC code.**

