## Supplementary Method 1. DLC code for "Next Generation AAV-F Capsid gene therapy rescues disease pathology in a model of Pyruvate Dehydrogenase Complex Deficiency"

Supplementary Methods

```
position_changes = {}

for group, dfs in group_data.items():
    position_changes[group] = {}

    for part in body_parts:
        change_list = []

        for df in dfs:
            # Check if both x and y columns for the body part exist
            if (part, 'x') in df.columns and (part, 'y') in df.columns:
                x_series = df[(part, 'x')]
                y_series = df[(part, 'y')]

                # Keep only valid (non-zero) x and y values
                valid_mask = (x_series != 0) & (y_series != 0)
                x_valid = x_series[valid_mask]
                y_valid = y_series[valid_mask]

                # Need at least 2 valid points to compute displacement
                if len(x_valid) >= 2 and len(y_valid) >= 2:
                    dx = np.diff(x_valid)
                    dy = np.diff(y_valid)
                    step_distances = np.sqrt(dx**2 + dy**2)
                    total_movement = np.sum(step_distances)
                    change_list.append(total_movement)
                else:
                    print(f"Not enough valid data for {part} in a file from group {group}.")
            else:
                print(f"Missing columns {(part, 'x')} or {(part, 'y')} in a file from group {group}.")

        if change_list:
            position_changes[group][part] = change_list
```
