## Supplementary Figures for "Next Generation AAV-F Capsid gene therapy rescues disease pathology in a model of Pyruvate Dehydrogenase Complex Deficiency"

### Supplementary Figure 1

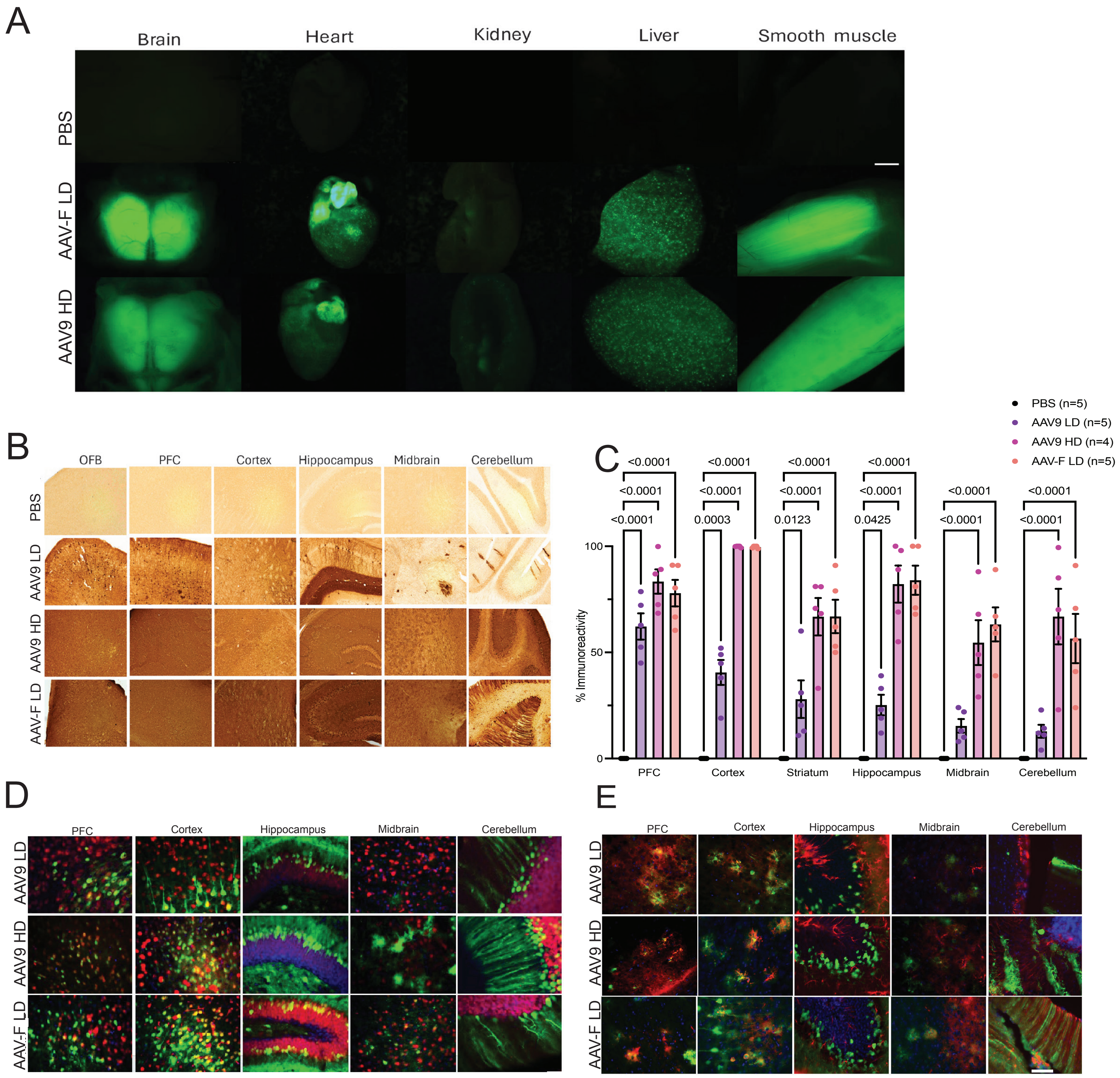

Supplementary Figure 2

A

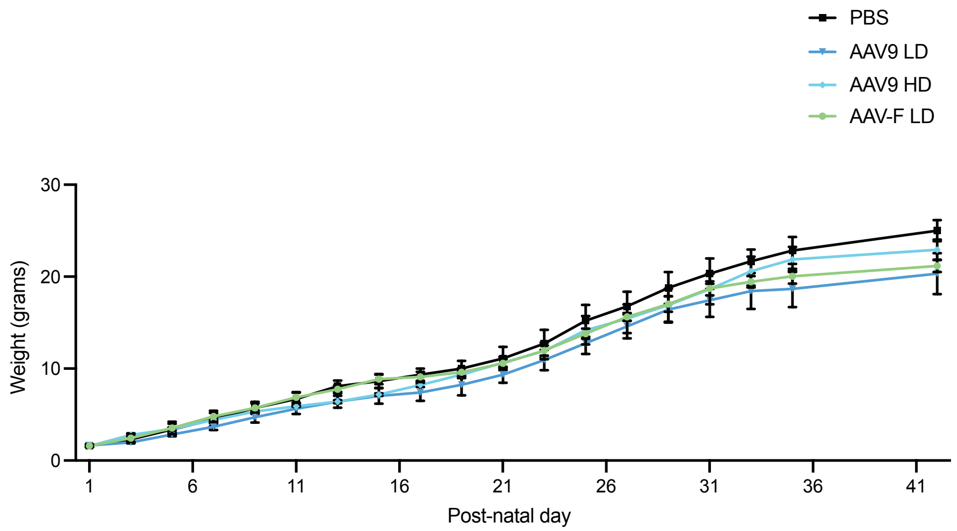


B

| Primer target | For/Rev/Probe | Sequence 5'-3' |
|---|---|---|
| *Pdha1* Genotyping | For | CGT CTG TTGAGA GAG CAG CA |
| *Pdha1* Genotyping | Rev | CGC ACA AGATAT CCA TTC CA |
| *hGFAP* Cre | For | ACT CCT TCATAA AGC CCT |
| *hGFAP* Cre | Rev | ATC ACT CGT TGC ATC GAC CG |
| *hPDHA1* | For | GGAGGATGGGCTCAAATACTAC |
| *hPDHA1* | Rev | GACCATCACACAAGTGACAGA |
| *hPDHA1* | Probe | ACGCCGAATGGAGTTGAAAGCAGA |
| *mPDHA1* | For | ATTCCTGGACTCAGGGTAGAT |
| *mPDHA1* | Rev | CCCTTACCAGACCTGCAATAG |
| *mPDHA1* | Probe | TATCTTGTGCGTCCGAGAGGCAAC |
| *mGapdh* | For | ACGGCAAATTCAACGGCAC |
| *mGapdh* | Rev | TAGTGGGGTCTCGCTCCTGG |
| *mGapdh* | Probe | TTGTCATCAACGGGAAGCCCATCA |

Supplementary Figure 3

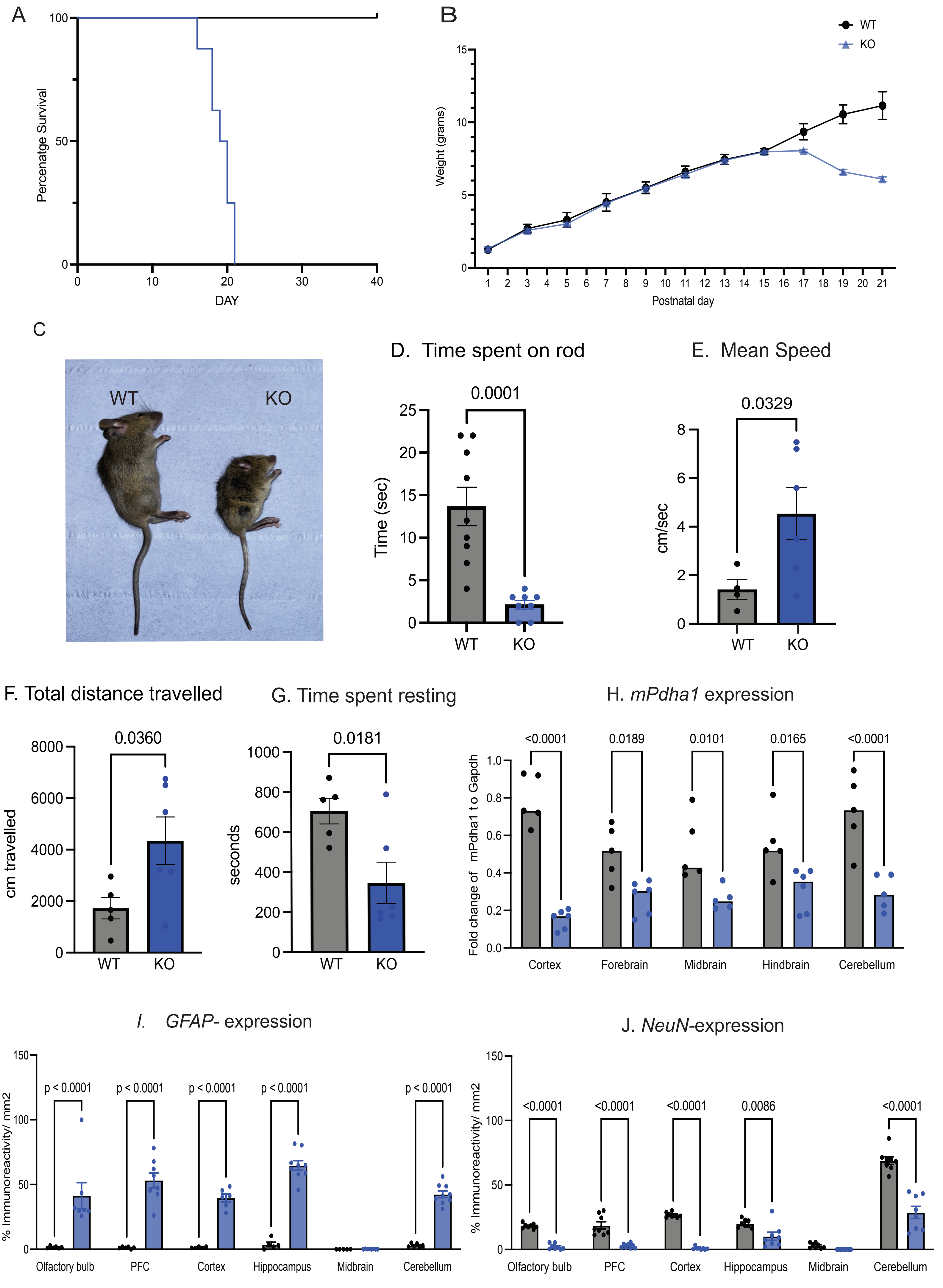

Supplementary Figure 4

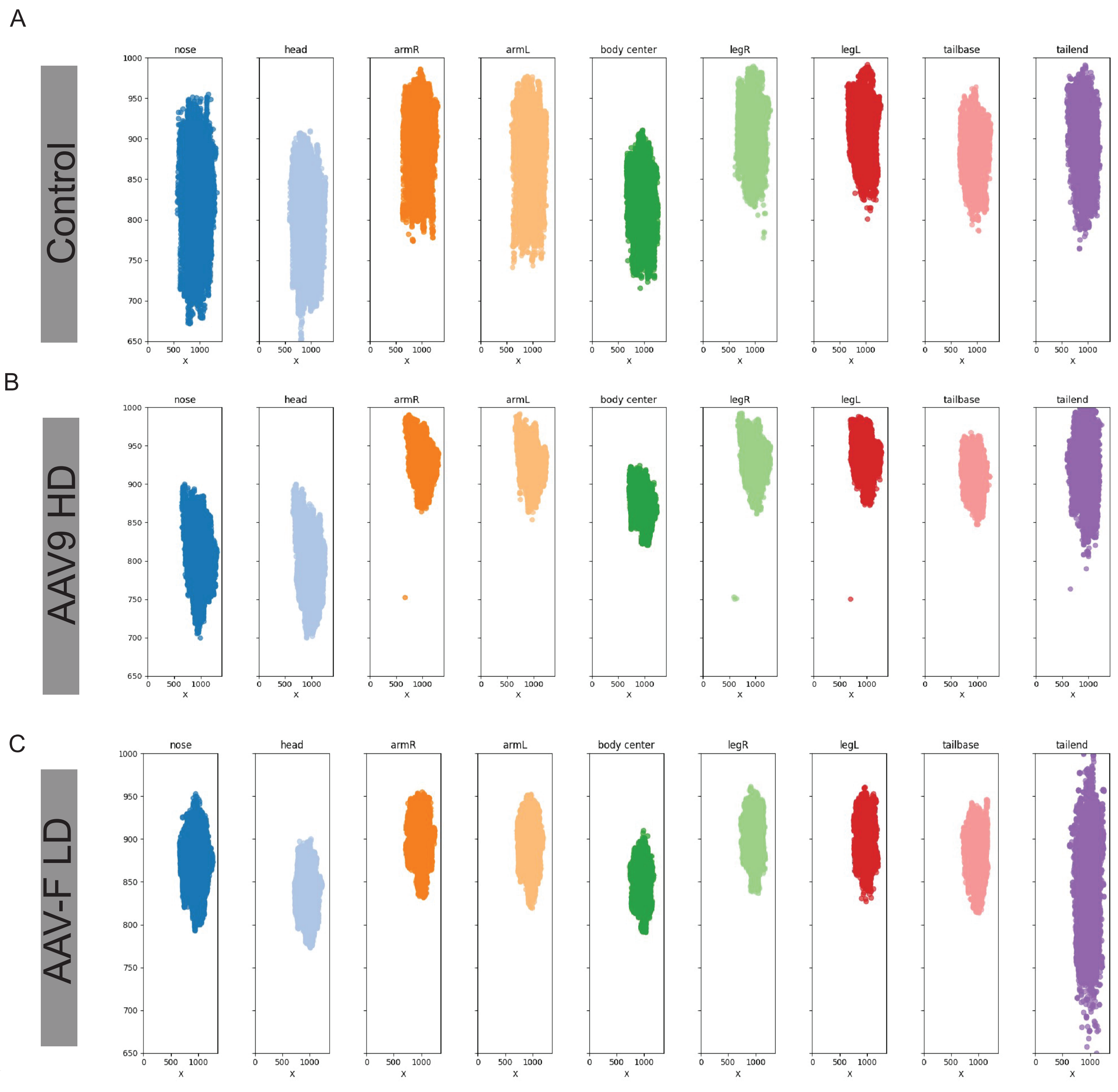

Supplementary Figure 5

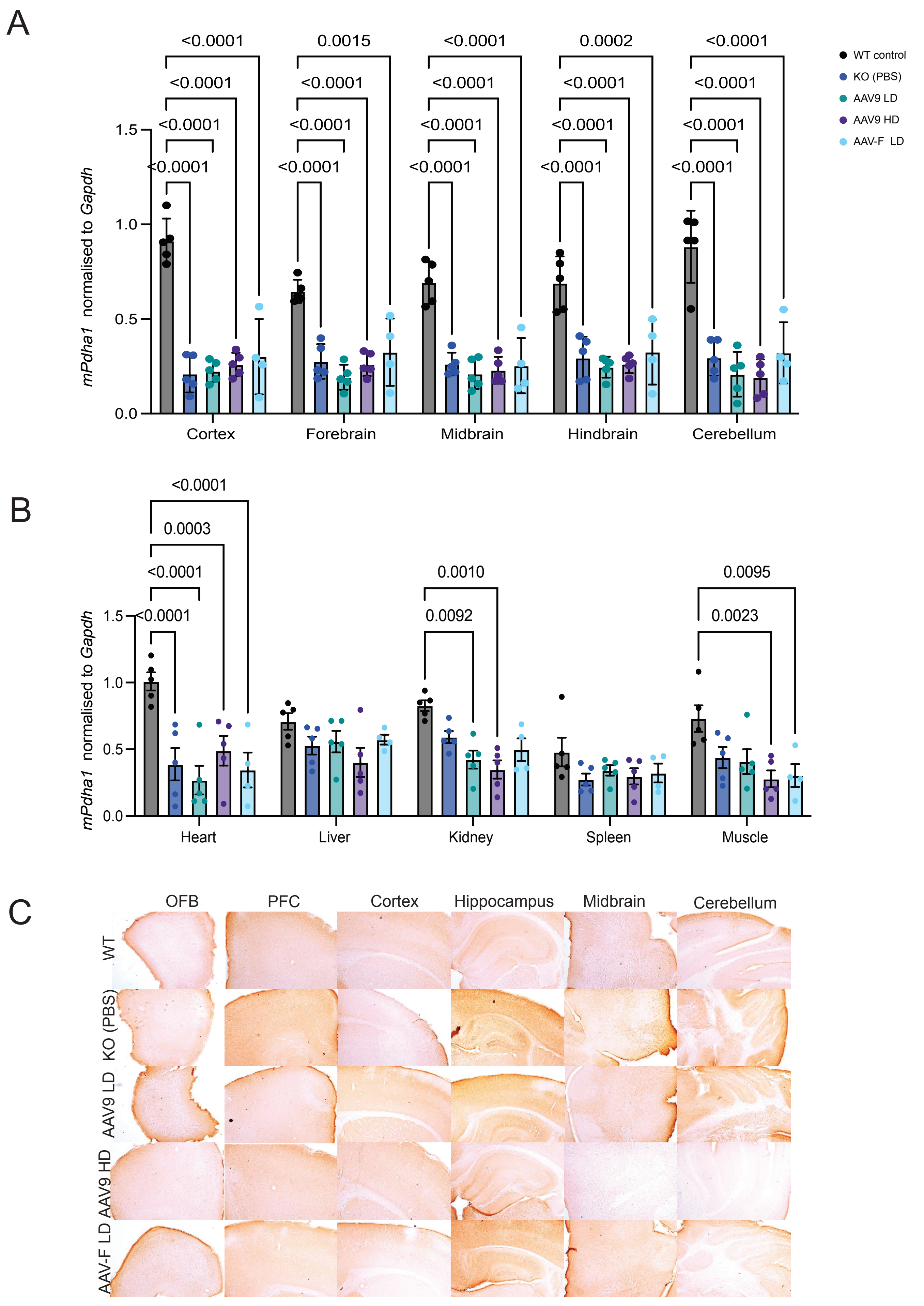
